# Shared hotspot mutations in spontaneously arising cancers position dog as an unparalleled comparative model for precision therapeutics

**DOI:** 10.1101/2021.10.22.465469

**Authors:** Lucas Rodrigues, Joshua Watson, Yuan Feng, Benjamin Lewis, Garrett Harvey, Gerald Post, Kate Megquier, Lindsay Lambert, Aubrey Miller, Christina Lopes, Shaying Zhao

## Abstract

Naturally occurring canine cancers have remarkable similarities to their human counterparts. In order to determine whether these similarities occur at the molecular level, we investigated hotspot mutations in a variety of spontaneously arising canine cancers and found high concordance in oncogenic drivers between cancers in both species. These findings suggest that canines may present a powerful and complementary model for preclinical investigations for targeted cancer therapeutics. Through analysis of 708 client-owned dogs from 96 breeds (plus mixed breeds) with 23 common tumor types, we discovered mutations in 50 well-established oncogenes and tumor suppressors, and compared them to those reported in human cancers. *TP53* is the most commonly mutated gene, detected in 30.81% of canine tumors overall and >40% in hemangiosarcoma and osteosarcoma. Canine tumors share mutational hotspots with human tumors in oncogenes including *PIK3CA, KRAS, NRAS, BRAF, KIT* and *EGFR*. Hotspot mutations with significant (P<0.0001) association to tumor type include NRAS G61R and PIK3CA H1047R in hemangiosarcoma, ERBB2 V659E in pulmonary carcinoma, and BRAF V588E in urothelial carcinoma. This work positions canines as excellent spontaneous models of human cancers that can help to investigate a wide spectrum of targeted therapies.

## Introduction

New approaches to the development of cancer therapeutics are urgently needed to improve the current 89% failure rate of novel drugs in clinical trials^1–3^ and improve patient outcomes. Spontaneous cancer in companion animals represents a unique opportunity for investigation of novel therapeutics for human and veterinary use^4–12^. Dogs develop spontaneous tumors which are highly similar to human cancers in histology and clinical presentation, but which often progress more quickly^7,10,11,13^. Canine cancers have also been found to have similar genetic and molecular targets to human malignancies^8,10,14–27^, and present an opportunity to test novel therapeutics in a treatment-naive setting, which is currently not feasible in human medicine^11^. Dogs represent a large animal model with an intact immune system, enabling comparative studies of therapeutic efficacy, immunotherapy, tumor evolution, and tumor microenvironment^7,11,28^. Because dogs also share our environment, they may share exposures to carcinogens, and could act as important environmental sentinels^29^. Studies of new therapeutic agents have begun to include dogs to help characterize pharmacokinetic and pharmacodynamic properties, efficacy, and tolerability^11,28^. In addition, the National Cancer Institute’s Center for Cancer Research has founded the Comparative Oncology Program and the Canine Oncology Trials Consortium to support comparative studies in dogs and facilitate integration of these findings with human oncology efforts^28^.

Canine tumors provide a powerful platform for translational investigation^11,12,30^. Over the past decade, genomic characterization of canine cancers has highlighted the close biological and molecular similarities between several canine and human cancers, including lymphoma^14,15,31^, osteosarcoma^8,20,22^, hemangiosarcoma^17,23–25,27,32^, glioma^26^, melanoma^19^, mammary tumors^33–35^, and urothelial carcinoma^16,18^. Some of the somatic mutations identified in these canine cancers occur at the orthologous position to known mutational hotspots found in human cancers, including PIK3CA H1047^17,23^, BRAF V588 (also reported as V595; human BRAF V600)^16^, and FBXW7 R470 (human R465)^15^.

Although studies have shown genomic concordance between canine and human cancers, the number of canine tumors sequenced via WES or WGS lags behind human tumors by an order of magnitude (fewer than 2000 canine tumors have been sequenced^36^, compared to more than 20,000 human tumors^37^). Consequently, the landscape of actionable tumor mutations in canine cancers is not fully understood^6,22^. We sought to address this issue in order to assess the feasibility of matching dogs with spontaneous cancers to targeted therapy, thereby providing treatment opportunities to canine patients while developing a platform that could accelerate a more global understanding of the clinical as well as translational potential for trial matching across species.

To do this, we developed a next-generation sequencing (NGS) panel targeting coding exons of 59 genes frequently mutated in human cancers. Using this panel, we performed the largest sequencing study of canine cancers to date, including 708 canine tumors from 23 tumor types from dogs representing more than 96 breeds. Importantly, our study revealed 20 canine mutational hotspots, 13 of which were orthologous to hotspots reported in human cancers, and clinically actionable. These results demonstrate significant overlap in somatic hotspot mutations between human and canine cancers, highlighting spontaneous canine cancers as an excellent model for the investigation of targeted therapies.

## Material and Methods

### Enrollment and sample collection

Client-owned dogs with histologically confirmed cancer diagnoses were enrolled in FidoCure® by 200 veterinarians in clinical practice. A total of 708 individual biopsies taken from May of 2019 until September of 2020 were analyzed through the FidoCure® Precision Medicine Platform, the proprietary name of The One Health Company’s precision medicine unit. Upon enrollment, tissue re-cuts obtained from formalin-fixed paraffin embedded (FFPE) tumor tissue used for histopathologic diagnosis were requested from the appropriate veterinary diagnostic laboratory. These tissues were evaluated by practicing board-certified veterinary pathologists and only tissue confirmed to be neoplastic progressed to genomic sequencing.

### Library Preparation and Next Generation Sequencing

Genomic DNA (gDNA) was extracted from FFPE tissues using Mag-Bind® FFPE DNA/RNA kit (Omega Bio-tek). The quality of the extracted gDNA was confirmed using the Agilent Genomic DNA ScreenTape Assay (Agilent) and the amount of gDNA was quantified using the Qubit dsDNA HS assay kit (Thermo Fisher). The DNA library was constructed using the SureSelect Low Input library prep kit (Agilent) according to the manufacturer’s protocol.

The FidoCure® Precision Medicine Platform targets the coding exons of the genes *ABL1, ALK, APC, ARID1A, ATM, BCL2, BCL6, BRAF, BRCA1, BRCA2, BTK, CDK2, CDK4, CDK6, CDKN2A, CREBBP, EGFR, ERBB2, FBXW7, FGFR1, FGFR2, FGFR3, FLT1/VEGFR1, FLT3, FLT4/VEGFR3, HDAC1, HIF1, HNF1, HRAS, JAK1, JAK2, JAK3, KDR/VEGFR2, KIT, KMT2C, KMT2D, KRAS, MEK/MAP2K1, MET, mTOR, NF1, NOTCH1, NRAS, TP53, PARP1, PDGFRa, PDGFRβ, PIK3CA, PTEN, PTPRD, PTPRT, RAF1, RB1, RET, ROS1, SETD2, SMAD4, SMARCA4*, and *TERT*. Genes commonly mutated in human cancers and those targeted by commercially available oncology panels were prioritized.

Hybrid capture-based enrichment of the targeted genes was performed using the SureSelect custom DNA Target Enrichment Probes and SureSelect XT Hyb and Wash kit following manufacturer’s instructions. The final library was quantified using qPCR and pooled for sequencing on the Illumina® platform (Illumina, California, USA) with a read length configuration of 150 PE for up to 6M PE reads (3M in each direction), yielding an approximate coverage of 612x per sample. Sequencing was performed in a CLIA-certified CAP-accredited laboratory.

### Variant calling and evaluation

The sequence read pairs were mapped to the canine reference genome (CanFam3.1) using BWA-MEM (v0.7.12)^38^. Mapped read coverage was obtained using GATK (version 3.8.1)^39^ AddOrReplaceReadGroup, MarkDuplicates, RealignerTargetCreator, and IndelRealgner, with minimum mapping quality of 10 and base quality of 10. BamTools^40^ was used to filter out low-quality reads (mapping quality < 5) and a high rate of mismatches (mismatch > 10).

A total of 1,181 unique variants were identified based on the pileup file prepared for each tumor sample by Samtools (version 1.9)^41^. Variants were also called using VarScan 2 (version 2.4.2)^42^, with default cutoffs (variants supported by at least three reads with base quality of ≥10; minimum variant allele fraction of 0.01, and minimum mapping quality of 20). Variants called by both methods were retained for further analysis. Variants were subjected to manual review and verification to identify errors including those in the canine gene annotation. Variant functional effects were annotated using SnpEff^43^ using the canFam31.99 annotation GTF file and SnpSift^44^. Synonymous, intronic, and untranslated region variants except for splice site variants were excluded. To reduce sequencing artifacts, variants with read coverage of ≤10, variant allele fraction (VAF) of <2%, or located in low-complexity regions (as defined by the RepeatMasker^45–48^ track on the UCSC Genome Browser) were also removed (Figure 2). Variant filtration was performed using Bcftools^49^ and SnpSift.

**Figure 1.**
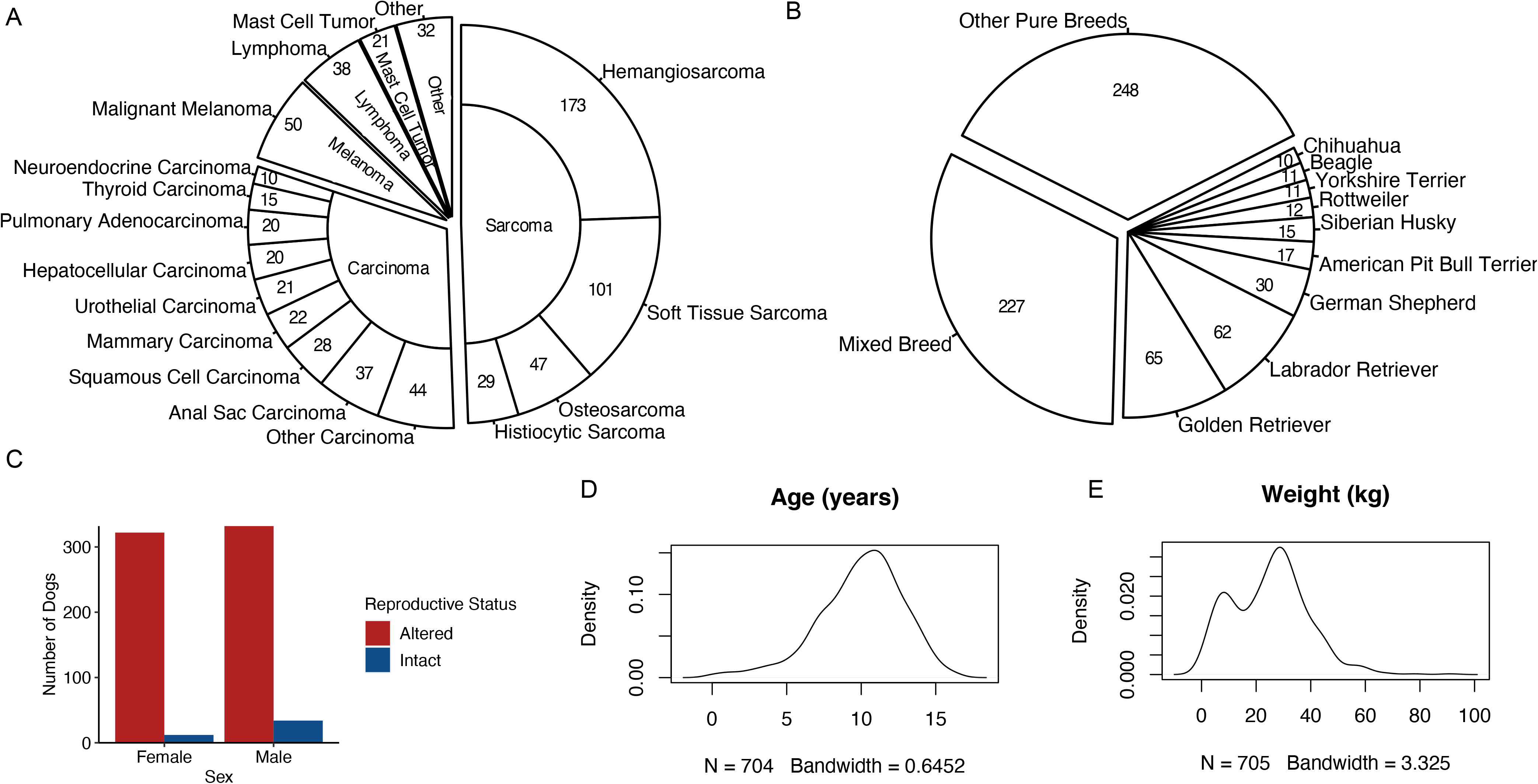
Demographics of enrolled dogs. A) Tumor types and supertypes. B) Breed (breeds with ≥10 dogs are identified).) C) Sex and reproductive status. D) Density plot of age. E) Density plot of weight.

**Figure 2.**
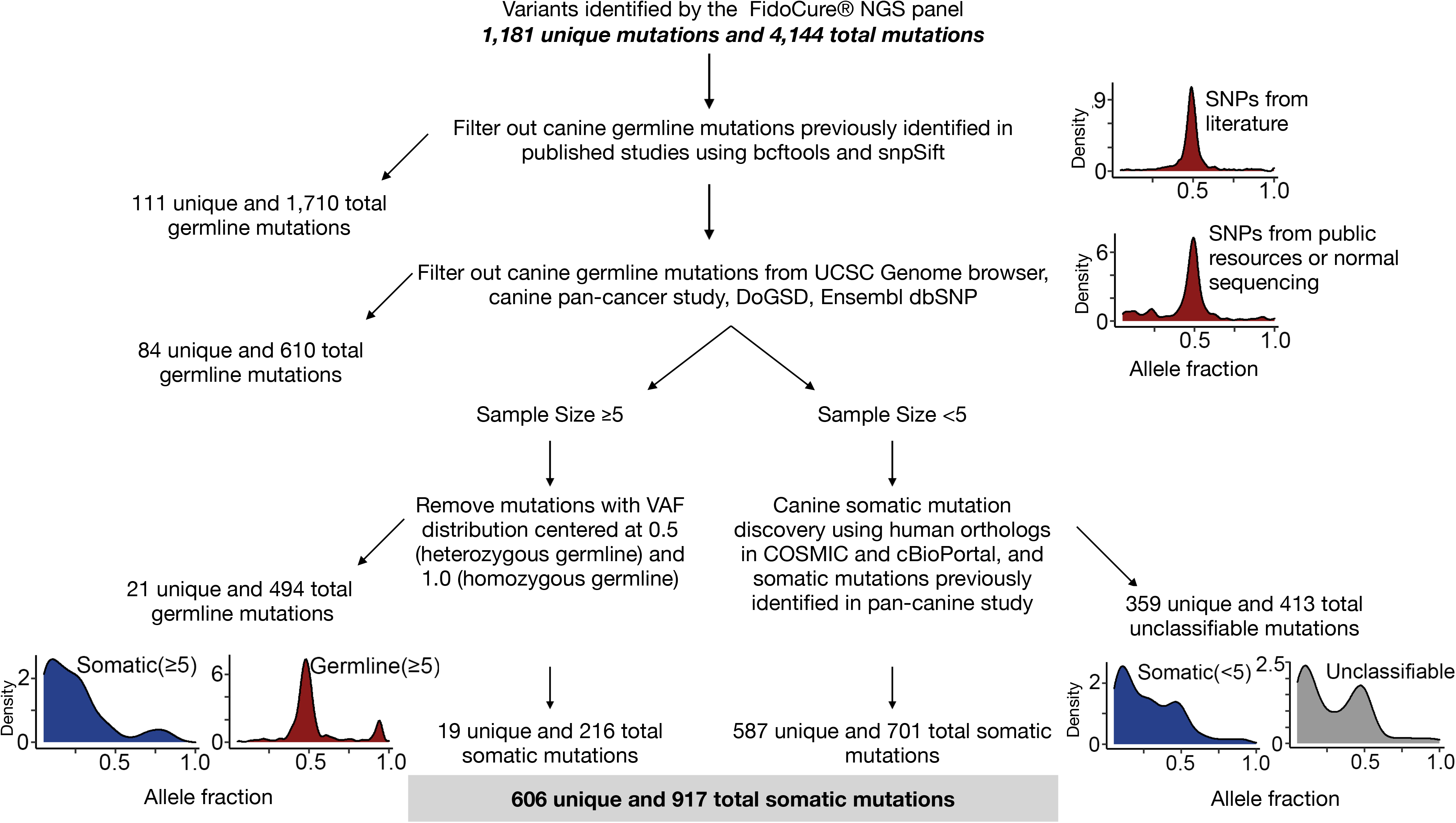
Germline-somatic mutation discrimination. Any variant already reported as germline mutation by previously published studies or public resources was excluded. Mutations identified in more than 5 dogs with variant allele fraction (VAF) distribution clustered at 0.5 (heterozygous germline) and near 100% (homozygous germline) were excluded. Mutations identified in fewer than 5 dogs were kept only if the orthologous human mutation was found in somatic mutation databases COSMIC or cBioPortal.

### Germline mutation filtering

As no matching germline samples from these dogs were sequenced, a subset of mutations were expected to be germline. Putative germline variants were identified and excluded in three steps as follows: 1) Variants with a VAF of >5% and total read count of >50 were compared to germline variants from >1,300 healthy dogs available in the European Variant Archive-EVA and other published studies^36,50^, Ensembl dbSNP^51^, the DoGSD website^52^, and 3,500,174 germline mutations from the Broad Improved Canine Annotation v1 track hub^53^ on the UCSC genome browser. 2) Variants found in ≥5 dogs with a VAF distribution clustered around 50% or near 100% were considered to be heterozygous or homozygous germline mutations and excluded. 3) For variants found in <5 dogs, the relevant human and canine protein sequences from the Ensembl annotations of the human (hg38, Release 103), and canine (CanFam3.1, Release 97) genomes were aligned using Clustal-Omega (version 1.2.4)^54^. The orthologous positions between the two species were identified by manual inspection of the protein alignments. Variants where the orthologous human mutation was reported in cBioPortal^55^ or the COSMIC database^56^ were classified as somatic. After filtering, 606 unique somatic mutational positions were identified.

### Definition of somatic mutational hotspots

Mutational hotspots in each species were annotated using the method developed by Chang, *et al*. ^57^, by identifying positions mutated more frequently than the background mutation with a cutoff of recurrence in >=4 samples (Supplementary Figure S2). Mutations at different nucleotide positions in the same codon of a gene and different nonsynonymous and synonymous base substitutions in the same codon were considered together.

### Statistical Analyses

Statistical analyses were performed using R (version 4.1.0)^58^. Fisher’s exact tests were used to compare mutation-positive and mutation-negative groups with categorical features to identify enrichment or depletion of variants in different categories. Multiple testing correction was applied using the Benjamini-Hochberg method. For all tests, a two-sided p-value of <0.05 was considered statistically significant. Enrichment scores were determined by -log10(q), with positive values indicating enrichment and negative values depletion.

## Results

### Cohort demographics

The final dataset consisted of somatic mutational data from 708 tumors from 23 cancer types. Hemangiosarcoma was the most common tumor type (n=173), followed by soft tissue sarcoma (n=101), melanoma (n=50), osteosarcoma (n=47), lymphoma (n=38) and anal sac carcinoma (n=37)(Figure 1A). In total, 350 sarcomas, 217 carcinomas, and 141 other cancer types were included (Figure 1A; Table S1). One dog had two samples from different cancer types submitted (anal sac carcinoma and soft tissue sarcoma), while all other enrolled dogs had one tumor sample submitted (Table S1).

The cohort included both purebred (n=481 dogs, 96 breeds), and mixed breed (n=227, ≥ 45 breeds) ancestry dogs (Table S1). A total of 9 breeds were represented by ≥10 dogs (Figure 1B; Table S1). The largest breed groups included golden retrievers (65 pure and 20 mixed), Labrador retrievers (62 pure and 30 mixed), German Shepherd Dogs (30 pure and 4 mixed) and American pit bull terriers (17 pure and 17 mixed) (Figure 1B). 370 cases were male (336 neutered and 34 intact) and 338 were female (325 spayed, 12 intact and 1 unknown) (Figure 1C; Table S1). Dogs ranged in age from 1 to 16 years (mean 9.9; SD 3.0) (Figure 1D, Table S1). Weight ranged from 1kg to 91kg (mean 24.0; SD 13.6) (Figure 1E; Table S1).

### Somatic mutations

The distributions of putative somatic and germline mutations among tumor types, breeds and age are consistent with published studies^15,17–20,22,23,26,36^. The base substitution distributions (Figure 4) are also consistent with those of known somatic and germline mutations identified in humans according to COSMIC, cBioPortal and Ensembl dbSNP.

**Figure 3.**
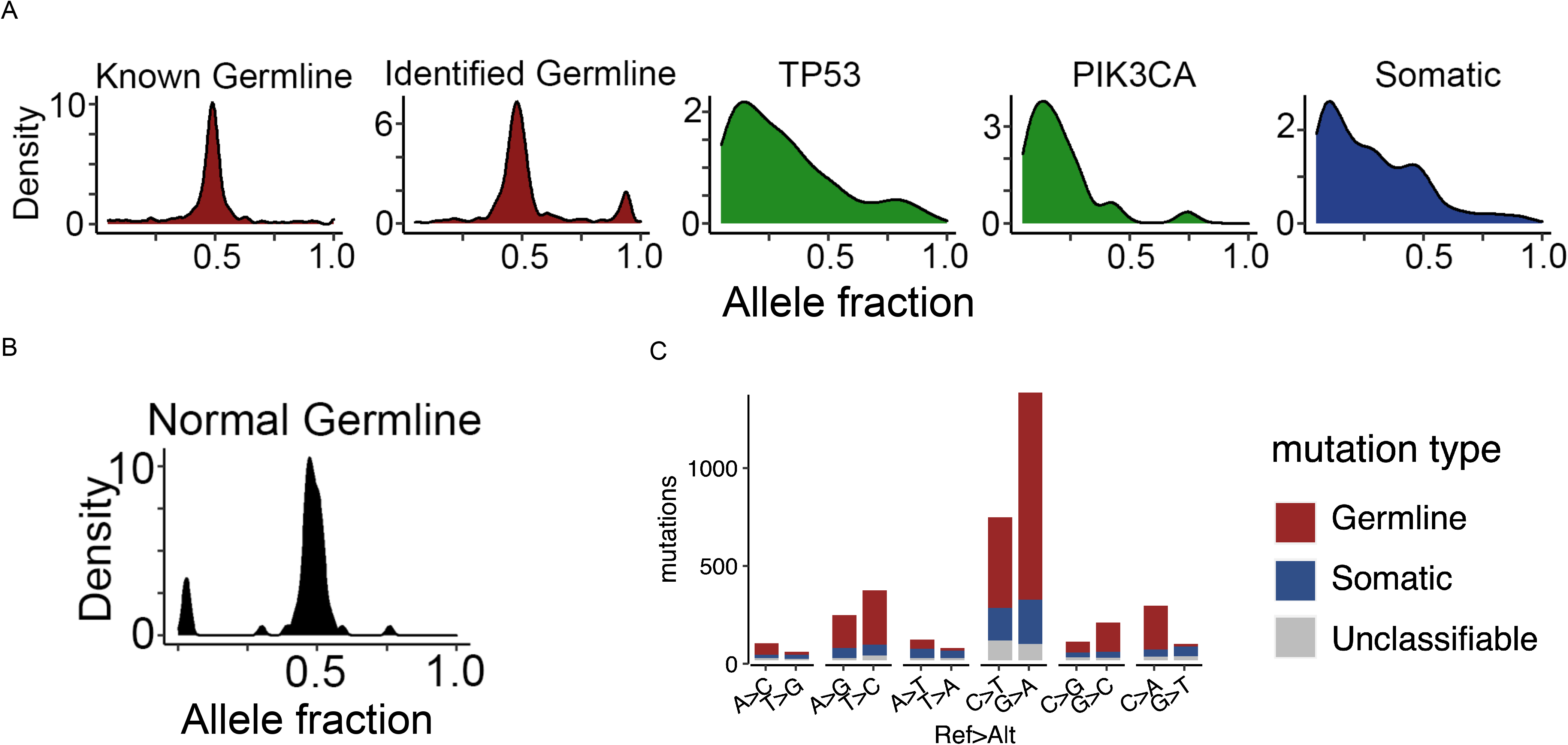
Identified germline-somatic mutation comparison. A) VAF distribution of germline mutations and somatic mutations identified from Figure 2. *TP53* and *PIK3CA* mutations are all somatic. B) VAF of 20 normal samples sequenced are also shown. C) Base substitution types: C>A, C>G, C>T and T>A, T>C and T>G distribution of somatic, germline and unclassified mutations.

**Figure 4.**
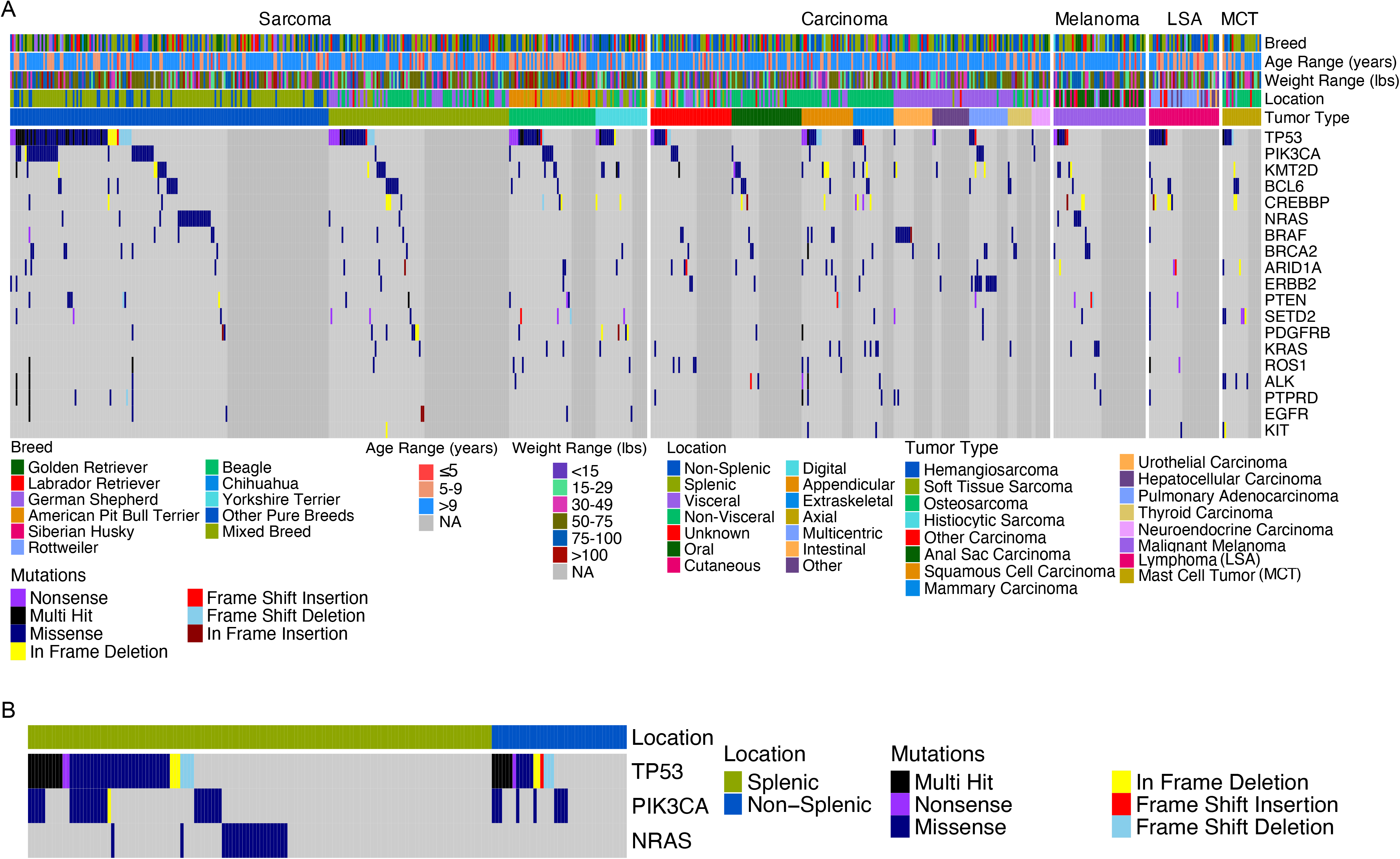
Somatic mutational landscape in canine tumors. A) Oncoprint of the 17 most frequently mutated genes in this data set and *EGFR* and *KIT* in each tumor sample, grouped by tumor supertype (sarcoma, carcinoma, melanoma, lymphoma, and mast cell tumor), breed, age, weight, and tumor location. B) Oncoprint showing mutually exclusive TP53, PIK3CA and NRAS mutations in splenic hemangiosarcoma.

We focused on putative somatic mutations (Table S2) for further analysis. *TP53* was the most frequently mutated gene (217/708, 30.6%) overall, as well as most frequently mutated in sarcoma, carcinoma, melanoma, and lymphoma (Figure 4). In 180 tumors, *TP53* carried a single mutation, while 35 tumors had two *TP53* mutations, and one tumor had three *TP53* mutations (Figure S1A). Other frequently mutated genes included chromatin remodelers (e.g., *KMT2D, SETD2, ARID1A, CREBBP*), PI3K signaling and MAPK signaling genes (e.g., *PIK3CA, PTEN, PDGFRB, ERBB2, BRAF, NRAS, KRAS*), and DNA repair genes (e.g., *BRCA2*) (Figures 4A and Figure S1A).

Recurrent mutational hotspots included PIK3CA His1047Arg (n=22), NRAS Gln61Arg (n=17), BRAF Val588Glu (n=14), ERBB2 Val659Glu (n=13), KMT2D Arg904Pro (n=13), NRAS Gln61Lys (n=8), TP53 Arg203Gly (n=8) (Figure S1B). These mutations are known or likely cancer drivers and most of them are activating or gain-of-function changes.

Our study provides a snapshot of somatic mutations for previously uncharacterized tumor types, including anal sac carcinoma and thyroid carcinoma. These two cancers have a significantly different mutational landscape than other types of carcinoma, including depletion of *TP53* mutations. Our study also provides a more comprehensive view of the mutational landscape for soft tissue sarcoma, hepatocellular carcinoma and mast cell tumor. We also observed a depletion of *PIK3CA* mutations for melanoma, lymphoma, mast cell tumors, urothelial carcinomas and thyroid carcinomas (Figure 4).

### Somatic mutation enrichment

We examined the distribution of recurrently mutated genes by tumor type and supertype. *TP53* mutations were significantly enriched in sarcoma (p= 4.2510^−6^), especially hemangiosarcoma (p= 8.83×10^−5^), but depleted in carcinoma (p= 8.6×10^−4^), especially anal sac carcinoma (p= 1.95×10^−3^). *PIK3CA* mutations were significantly enriched in hemangiosarcoma (p= 1.88×10^−7^), (Figure 5A). *NRAS* mutations were enriched in hemangiosarcoma (p= 2.27×10^−5^), and were mutually exclusive with *TP53* (P<0.01) and *PIK3CA* (P<0.05) mutations in splenic hemangiosarcoma (Figure 4B), as well as being depleted in carcinomas. Other significant enrichments included *ERBB2* mutations in pulmonary adenocarcinoma (p= 4.71×10^−11^) and *BRAF* mutations in urothelial carcinoma (p= 2.98×10^−6^).

**Figure 5.**
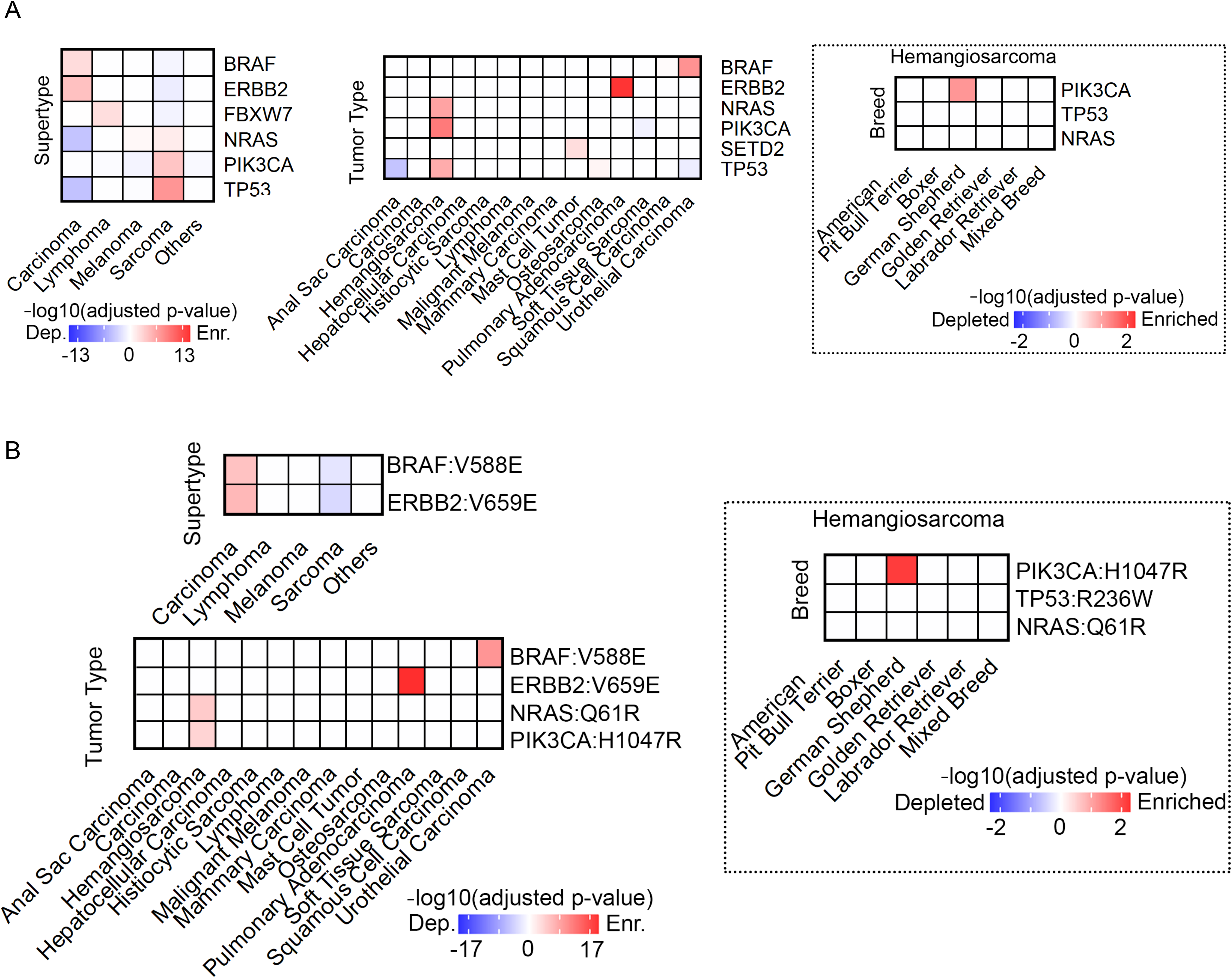
Canine somatic mutation enrichment and association. Shown are heatmaps indicating the enrichment (red) or depletion (blue) scores (based on Fisher’s exact test) of (A) genes mutated in >5 dogs and (B) individual mutations present in >5 dogs, grouped by tumor supertype, tumor type (with >20 samples), and breed.

We also examined the distribution of recurrent individual mutations (Figure 5B). *BRAF* Val588Glu and *ERBB2* Val659Glu are significantly enriched in carcinomas, but depleted in sarcomas (Figure 5B). Significant enrichment among tumor types include *BRAF* Val588Glu in pulmonary adenocarcinoma and *BRAF* Val588Glu mutation in urothelial carcinoma (Figure 5B).

We assessed whether any somatic mutations were associated with breed background. PIK3CA His1047Arg mutations were enriched in German Shepherd Dogs (p= 0.03) (Figure 5A-B). However, dogs with German Shepherd ancestry in this cohort were likely to have hemangiosarcoma (24/35, 69%), which also had a strong association with PIK3CA His1047Arg mutations. No other significant enrichment or depletion was detected within breeds.

### Comparative analysis of mutational hotspots

We identified 20 canine mutational hotspots, which are more likely to be cancer drivers^57,59^ and anti-cancer targets (Supplementary Table S3). Many of the mutational hotspots identified were in oncogenes, including *PIK3CA, KRAS, NRAS, BRAF, KIT, ERBB2*, and *EGFR* (Figure 6). Mutational hotspots were also identified in *TP53*, a tumor suppressor.

**Figure 6.**
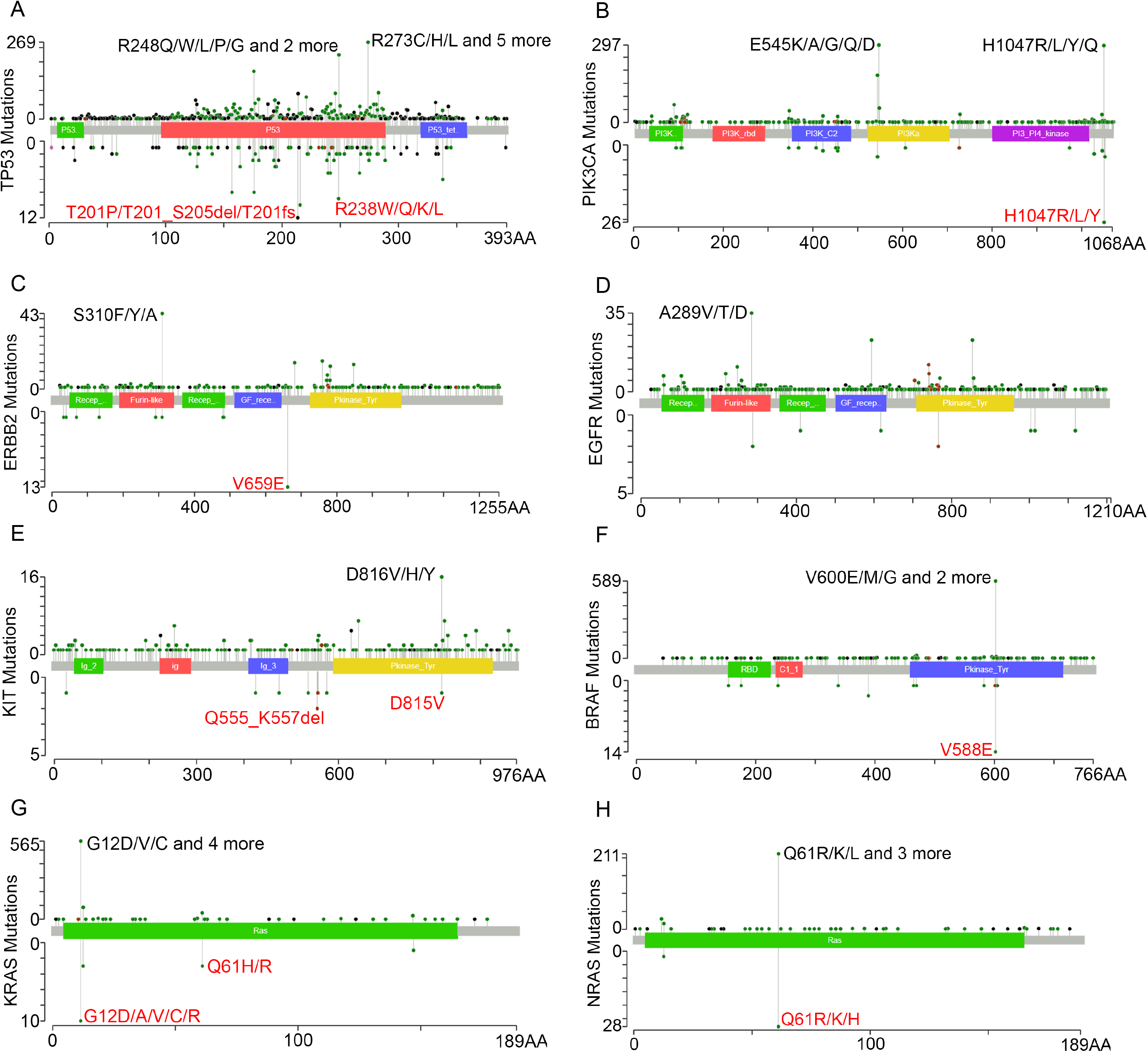
Comparison of canine and human mutational hotspots. Lollipop plots depict the mutational distribution in (A) TP53, (B) PIK3CA, (C) ERBB2, (D) EGFR, (E) KIT, (F) BRAF, (G) KRAS, and (H) NRAS in 24,592 human (upper) and 708 canine (lower) tumors. Amino acid position in the human protein is shown on the x-axis. Number of mutations is shown on the y-axis. The most prominent mutational hotspots are labeled, with the canine position also indicated.

We compared these canine mutational hotspots to those reported in 24,592 human tumors^57,60^. Many human mutational hotspots such as PIK3CA H1047, KRAS G12, NRAS Q61, and BRAF V600 (canine V588) are also seen in the orthologous amino acid position in dogs. In addition, species-specific mutational hotspots were identified, including PIK3CA E542/E545 and ERBB2 S310 in humans, as well as ERBB2 V659 in dogs (Figure 6).

The majority of mutations identified in KRAS were located at amino acid position 12, with mutations including G12D (2 mammary carcinomas, 1 carcinoma, and 1 squamous cell carcinoma), G12V (2 mammary carcinomas), G12A (1 melanoma), G12C (1 melanoma), and G12R (1 squamous cell carcinoma). In *NRAS* (which has 100% homology in the amino acid sequence between dogs and humans) G12 and Q61 were the most common positions mutated. We identified the Q61 mutation in 30 canine tumors, 70% (21/30) of which were hemangiosarcoma, 17% (5/30) malignant melanoma, 7% (2/30) plasma cell tumors and 7% (2/30) soft tissue sarcoma.

## Discussion

We performed a mutational hotspot analysis of 708 spontaneous canine tumors across 23 tumor types and 96 breeds common to pet dogs in the US. To our knowledge, this represents the largest sequencing study of canine tumors to date, and includes tumor types whose mutations have not previously been characterized (*e*.*g*., anal sac carcinoma, thyroid carcinoma and neuroendocrine carcinoma) or have been less characterized (*e*.*g*., soft tissue sarcoma, hepatocellular carcinoma and mast cell tumor). To do this, we developed the FidoCure® Personalized Genomic Panel, a targeted sequencing panel containing 59 well-known oncogenes and tumor suppressors frequently mutated in human cancer, with common mutational hotspots. Our study adds to the growing body of canine comparative oncology studies demonstrating genomic similarities between human and canine cancers, and specifically evaluates hotspot mutations that can be targeted with a precision medicine approach.

Our analysis largely captures the landscape of hotspot mutations in canine tumors, which were similar to the mutational landscapes reported by previous whole exome or genome sequencing studies (e.g., *TP53, NRAS* and *PIK3CA* in hemangiosarcoma, *TP53* and *SETD2* in osteosarcoma, *ERBB2* in pulmonary carcinoma, *BRAF* in urothelial carcinoma and *FBXW7* in lymphoma)^15,17,19–24,26,33–36,61,62^. Consistent with previous research, our results indicate that *TP53* is the most recurrently mutated gene across tumor types^15,17,19–24,26,33–36,61,62^. *TP53* mutations were significantly more common in sarcomas (p= 2.2×10^−6^) than in carcinomas (p= 4.7×10^−4^). This may be due to differences in the cell of origin and the mechanisms of tumorigenesis in these cancer types. Carcinomas originate from polarized epithelial cells or their progenitors, and alterations of cell polarity genes and loss of cell polarity are likely the major drivers of accelerated cell proliferation in carcinoma development^63–65^. Sarcomas originate from mesenchymal cells, and loss of function of *TP53*, which leads to defective cell cycle checkpoints and accelerated proliferation. Our studies also identified other frequently mutated genes reported in canine cancer, including PI3K signaling genes (*PIK3CA*), RAS/MAPK signaling genes (*NRAS* and *KRAS*), and chromatin modeling genes (*KMT2D* and others).

Mutational landscape varied by tumor type, but was largely breed-independent, consistent with a previous pan-cancer and pan-breed study that investigated previously published whole exome data from 591 canine tumors^36^. The largest difference was between carcinomas and sarcomas, with significant differences in mutational frequency for *TP53, PIK3CA, NRAS, ERBB2* and *BRAF*. Carcinomas were more variable than sarcomas in their mutational spectrum. Anal sac carcinoma, thyroid carcinoma and neuroendocrine carcinoma, for which sequencing data has not previously been published, were significantly different from other carcinomas, and were depleted for *TP53* mutations. While German Shepherd Dogs were more likely to have a *PIK3CA* His1047Arg mutation than other breeds, they were also more likely to have hemangiosarcoma, which was enriched for the *PIK3CA* His1047Arg mutation.

We found that *NRAS* mutations were mutually exclusive with *TP53* and *PIK3CA* mutations in splenic hemangiosarcoma, reaffirming the existence of different molecular subtypes of the same histology type^24^.

Our work reveals that numerous mutational hotspots are shared between dogs and humans, including *PIK3CA* H1047, *BRAF* V600/V588, *KRAS* G12 and others. These findings further position dogs as a powerful translational model for human and veterinary oncology, as both existing and novel targeted therapies for these mutations (*e*.*g*. PIQRAY for *PIK3CA* H1047R mutations^66^, PLX4032 for *BRAF* V600E mutations^67^) can be assessed in canine cancer patients.

Our analysis also revealed species-specific mutational hotspots, including *PIK3CA* E545/2K mutations found only in human cancers. Further studies will be needed to better understand the mechanisms underlying these differences, which will assist anti-cancer drug development and precision medicine in both species.

The FidoCure® Personalized Genomic Panel effectively captures the landscape of actionable hotspot mutations in canine tumors. We anticipate that this resource will help to accelerate canine cancer genomic research, significantly increasing the use of the canine model in precision medicine and anti-cancer drug development both as a biomedical model and to benefit veterinary patients.

## Acknowledgments

This study was conducted by The One Health Company and its FidoCure® Partners. Work performed at UGA is supported by the National Cancer Institute grants R01 CA182093 and CA252713. We deeply thank the pet owners who entered their dogs in our study during such a painful time, and all of the veterinarians who treated these amazing dogs: Aarti Sabhlok, Adam Breiteneicher, Alexandra Barrientos, Alice Villalobos, Alicia Andras, Alison Book, Alyssa Murray, Amanda Beck, Amanda Elpiner, Amanda Foskett, Amanda Shively, Amanda Smith, Amber Tegtmeyer, Amelia Keith, Amy Back, Amy Nichelason, Andi Flory, Andrew Scarborough, Angela Taylor, Angie Stamm, Anthony Gray, Ariana Verrilli, Arnold Lesser, Ashley Davis, Audrey Stevens, Aurélia Klajer, Avenelle Turner, Barbara Kitchell, Ben Spitz, Ben Staiger, Benjamin Lee, Beth Overley-Adamso, Betsy Hershey, Birgitte Tan, Bisque Jackson, Blaise Burke, Bonnie Smith, Brenda Phillips, Brenda Tam, Brennen McKenzie, Bret Hixson, Brian Butzer, Bridget Urie, Brolin Evans, Brooke Britton, Brooke Fowler, Caleb Alexander, Carlos Canino, Carlos Rodriguez Jr, Carly Campbell, Carrie Donahue, Carrie Uehlein, Carrissa Wood, Catriona D’Aulerio, Charles Wilkins, Chelsea Davis, Christina Choate, Christina Manley, Christine Mullin, Christine Oakley, Christine Swanson, Cindy Bressler, Colleen Martin, Colleen Tansey, Craig Clifford, Dan Polidoro, David Bommarito, David Glover, David Heller, David Hunley, Dean Vicksman, Deborah Knapp, Debra Canapp, Diane Cosko, Diane Craig, Diane Schrempp, Dorothy Jackson, Douglas Bahr, Drew Humphries, Edith Karapet, Edmund Sullivan, Ednee Yoshioka, Edwina Love, Eileen Savier, Elizabeth Schuh, Emily Wilson, Emma Katz, Eric Bulakowski, Erica Faulhaber, Francisco Alvarez, Frank Defeis, Gene Solomon, Gerald Post, Gina Kwong, Gregory Bogatsky, Haley Williams, Heather Reeder, Heidi Ward, Holly Burr, Ian Muldowney, Jaclyn Renner, Jacqueline Wypij, Jamie Wignall, Jana Zapalac, Janet Peterson, Jarred Lyons, Jeff Philibert, Jennifer Baez, Jennifer Danielson, Jennifer Hauss, Jennifer Hofer, Jennifer McDaniel, Jennifer Pierro, Jennifer Wiley, Jessica Chin, Jessica Miller, Jessica O’Neill, Jessica Thiman, Ji-In Lee, Jim Perry, John Reed, Joseph Impellizeri, Joshua Lachowicz, Julie Bulman-Fleming, Kara Magee, Karen Johnston, Karina Valerius, Karri Miller, Kate Raczak, Katherine Wright, Kathryn Beiser, Katie Wright, Kelley Cox, Kendra Lyons, Kerry Rissetto, Kim Cronin, Kimberly Alexander-Coleman, Kimberly Freeman, Kimberly Menard, Kirkwood Vardeman, Klarissa Ligon, Krista Seraydar, Krystal Harris, Lauren Blaeser, Lauren Sikorski, Lawrence Putter, Lindsay Thalheim, Lindsey Fry, Lisa Fulton, Lisa Knapp, Lisa Parshley, Loren Nations, Luminita Sarbu, Lynda Beaver, Lyndsay Kubicek, Marc Wallach, Margret Puccio, Marina Tejada, Mark Anderson, Mark Moore, Mary Davis, Mary Kay Blake, Matthew Sherger, Maura Faller, Megan Duckett, Megan Howard, Meighan DeHart, Melina Maritato, Meline Joaris, Melissa Miller, Melissa Parsons-Doherty, Meredith Gauthier, Merrianne Burtch, Michael Dixon, Michael Kiselow, Michael Linderman, Michele Cohen, Michelle Morges, Mike Pallone, Milinda Lommer, Mona Rosenberg, Nancy Currin, Naoko Sogame, Natalie Goldberger, Nathaniel Vos, Nicola Wilson, Nicole Leibman, Nirip Shokar, Noelle Bergman, Oceane Aubry, Pamela Lucas, Patrick Cutbirth, Paul Berdoulay, Paul Morgan, Paula Bratich, Pavel Mihok, Pedro Dominguez, Rachel Galluzzo, Rachel Reiman, Rachel St-Vincent, Rachel Venable, Ravinder Atwal, Rebecca Brown, Rebecca Kilcullen, Rebecca Newman, Rhonda Feinmehl, Rick Wassell, Rita Ho, Robert Proietto, Robert Swinger, Robert Woods, Roberta Portela, Ron Weich, Saber Mansour, Sally Gilbertson, Samantha Bajorek, Sara Fiocchi, Sara Fritz, Seth Glasser, Shana Hutchison, Shana O’Marra, Sharon Nath, Sherri Sprague, Shirley Chu, Sindy Piscoya, Stacy Binstock, Stacy McLeod, Stephen Atwater, Stephen Brenn, Steve Larson, Sue Downing, Susan Lana, Susan Sundburg, Susie Kang, Takashi Kasuya, Tanya Gustafson, Terrance Hamilton, Theresa Arteaga, Timothy Estabrooks, Todd Erfourth, Tonya Curtis, Traci Lazar, Trina Hazzah, Tristram Bennett, Valerie Wiles, Victor Brown, Vincent Baldanza, Virginia Phelps, Wendy Ellis, and Wendy Lavalle.

